# Role of ACSBG1 in brain lipid metabolism and X-linked adrenoleukodystrophy pathogenesis: Insights from a knockout mouse model

**DOI:** 10.1101/2024.06.19.599741

**Authors:** Xiaoli Ye, Yuanyuan Li, Domingo González-Lamuño, Zhengtong Pei, Ann B. Moser, Kirby D. Smith, Paul A. Watkins

## Abstract

The "bubblegum" acyl-CoA synthetase (ACSBG1) is a pivotal player in lipid metabolism during the development of the mouse brain, facilitating the activation of long-chain fatty acids (LCFAs) and their integration into essential lipid species crucial for brain function. Through its enzymatic activity, ACSBG1 converts LCFAs into acyl-CoA derivatives, supporting vital processes like membrane formation, myelination, and energy production. Its regulatory role significantly influences neuronal growth, synaptic plasticity, and overall brain development, highlighting its importance in maintaining lipid homeostasis and proper brain function.

Originally discovered in the fruit fly brain, ACSBG1 attracted attention for its potential implication in X-linked adrenoleukodystrophy (XALD) pathogenesis. Studies using Drosophila melanogaster lacking the ACSBG1 homolog, bubblegum, revealed adult neurodegeneration with elevated levels of very long-chain fatty acids (VLCFA). To explore ACSBG1’s role in fatty acid (FA) metabolism and its relevance to XALD, we created an ACSBG1 knockout (Acsbg1^-/-^) mouse model and examined its impact on lipid metabolism during mouse brain development.

Phenotypically, Acsbg1^-/-^ mice resembled wild type (w.t.) mice. Despite its primary expression in tissues affected by XALD, brain, adrenal gland and testis, ACSBG1 depletion did not significantly reduce total ACS enzyme activity in these tissues when using LCFA or VLCFA as substrates. However, analysis unveiled intriguing developmental and compositional changes in FA levels associated with ACSBG1 deficiency.

In the adult mouse brain, ACSBG1 expression peaked in the cerebellum, with lower levels observed in other brain regions. Developmentally, ACSBG1 expression in the cerebellum was initially low during the first week of life but increased dramatically thereafter.

Cerebellar FA levels were assessed in both w.t. and Acsbg1^-/-^ mouse brains throughout development, revealing notable differences. While saturated VLCFA levels were typically high in XALD tissues and in fruit flies lacking ACSBG1, cerebella from Acsbg1^-/-^ mice displayed lower saturated VLCFA levels, especially after about 8 days of age. Additionally, monounsaturated ω9 FA levels exhibited a similar trend as saturated VLCFA, ω3 polyunsaturated FA levels werewhile elevated in Acsbg1^-/-^ mice.

Further analysis of specific FA levels provided additional insights into potential roles for ACSBG1. Notably, the decreased VLCFA levels in Acsbg1^-/-^ mice primarily stemmed from changes in C24:0 and C26:0, while reduced ω9 FA levels were mainly observed in C18:1 and C24:1. ACSBG1 depletion had minimal effects on saturated long-chain FA or ω6 polyunsaturated FA levels but led to significant increases in specific ω3 FA, such as C20:5 and C22:5.

Moreover, the impact of ACSBG1 deficiency on the developmental expression of several cerebellar FA metabolism enzymes, including those required for synthesis of ω3 polyunsaturated FA, was assessed; these FA can potentially be converted into bioactive signaling molecules like eicosanoids and docosanoids.

In conclusion, despite compelling circumstantial evidence, it is unlikely that ACSBG1 directly contributes to the pathology of XALD. Instead, the effects of ACSBG1 knockout on processes regulated by eicosanoids and/or docosanoids should be further investigated.

## INTRODUCTION

Fatty acids (FA) are the building blocks of complex lipids, including triacylglycerol, glycerophospholipids, sphingolipids and cholesterol esters. Fatty acids are also an indispensable metabolic fuel when degraded by β-oxidation. To participate in either anabolic or catabolic pathways, fatty acids must first be activated by thioesterification to coenzyme A (CoA), a reaction catalyzed by members of the fatty acyl-CoA synthetase (ACS) family (EC 3.4.1.x) [1,2]. Phylogenetic analysis revealed that most human and mouse ACSs segregate into subfamilies that roughly correlate with their FA substrate chain length preference; thus, short- (ACSS), medium- (ACSM), long- (ACSL), and very long-chain (ACSVL) ACSs have been described [2]. Human and mouse homologs of the gene disrupted in the Drosophila melanogaster “*bubblegum*” mutant was the basis for identification of an additional ACS family, designated ACSBG [3]. In addition to differences in FA chain-length preference, ACSs also differ in their tissue, cell, and subcellular locations [1]. Most tissues and cells express several ACSs. For example, proteomics indicated that brain astrocytes express at least 7 of the 14 ACSs that constitute the ACSL, ACSVL, and ACSBG families [PAW, unpublished observation]. These observations suggest that individual ACSs must play unique and specific roles in lipid metabolism.

Several inherited neurologic disorders, particularly the leukodystrophies, are associated with abnormal FA metabolism. In X-linked adrenoleukodystrophy (XALD), deficient degradation of saturated very long-chain FAs (VLCFA) in peroxisomes results in elevated levels of these FAs, and in particular C26:0, in plasma and tissues [4]. Early hypotheses predicted defective synthesis of VLCFA-CoA by a peroxisomal very long-chain ACS was the cause of XALD [5,6]. Discovery of the gene mutated in XALD, ABCD1, largely disproved this hypothesis [7]. ABCD1 is an ATP-binding cassette half-transporter and is predicted to homodimerize in the peroxisomal membrane to form a functional transport molecule [8]. Subsequent investigation revealed that ABCD1 does not transport VLCFA, but rather VLCFA-CoA [9–11].

There are two major phenotypic presentations of XALD – the childhood cerebral form (CCER) and the adult-onset peripheral neuropathy, adrenomyeloneuropathy (AMN) [4]. Both CCER and AMN are caused by mutations in ABCD1, and both phenotypes can often be present in members of the same nuclear family. In CCER, which affects about 35% of patients, symptoms typically appear around 7 years of age. Inflammatory demyelination usually leads to death by 4 years following onset of cerebral symptoms. In AMN, which affects more than 60 % of patients, the central nervous system is generally spared, but a dying back of long axons of spinal cord neurons leads to a disturbance of gait as well as effects on bowel and blader function. Almost all XALD patients have adrenal insufficiency (Addison’s disease), and this typically predates neurologic symptoms. Testicular insufficiency is frequently seen in AMN patients. Thus, the primary organs affected in XALD are brain and nerve, adrenal glands, and testis. Since ABCD1 is expressed in most tissues (Suppl. Fig. 1), the question arises as to why mutations have pathologic consequences in only a few tissues. Of potential significance is the fact that brain expression of ABCD1 is among the lowest of 27 tissues examined, while adrenal and testis expression are among the highest (Suppl. Fig. 1).

Although abnormal activation of VLCFA is thus not causative of XALD pathology, the nature of the specific ACS that activates these FA prior to transport, particularly in the tissues pathologically affected in this disorder, has not been determined. Among the ACSs that are potential candidate activators of VLCFA, one enzyme of potential interest is ACSBG1. This enzyme has been characterized in humans [12,13] and mice [14–15], and is also expressed in many other organisms [16].

We [12,13] and others [14] reported that ACSBG1 expression is primarily found in brain, adrenal gland, testis, and ovary. Deficiency of the ACSBG1 homolog, *bubblegum*, in Drosophila melanogaster led to adult neurodegeneration with marked dilation of photoreceptor axons and elevated levels of VLCFA [3]. To gain further insight into the potential role of ACSBG1 in FA metabolism, and to assess any potential relevance to XALD, we produced an ACSBG1 knockout (KO) mouse. In this report, we examine the potential role of ACSBG1 in lipid metabolism in the developing mouse brain.

## MATERIALS AND METHODS

### Materials and General Methods

Affinity-purified antibody to human ACSBG1 was prepared as described previously [13]. Sources of commercial primary antibodies were as follows: GAPDH (Sigma-Aldrich), ACC1 (Cell Signaling), FASN (Cell Signaling). Secondary antibodies were from Jackson ImmunoResearch. Protein was measured using the Pierce 660 nm protein assay kit (Thermo scientific, USA) according to the manufacturer’s protocol. [1-14C]palmitic acid (C16:0) and [1-14C]lignoceric acid (C24:0) were from Moravek Inc. ACS activity in mouse tissues was measured as described previously [17]. RNA-seq data for estimation of ABCD1 expression in 27 human tissues was obtained through the National Center for Biotechnology Information (NCBI) portal [18] and is shown in Suppl. Fig. 1.

### Animals and their care

Mice (strain 129vev) were obtained from Taconic Biosciences. All animal studies were approved by the Johns Hopkins University School of Medicine Institutional Animal Care and Use Committee (IACUC) in accordance with the guidelines and regulations described in the NIH Guide for the Care and Use of Laboratory Animal. Wild type (w.t.) and Acsbg1 knockout mice (Acsbg1^-/-^) were housed in the animal facility of Johns Hopkins School of Medicine on a 12-hour light/dark cycle with ad libitum access to food and water. The facility is pathogen-free and is maintained at a constant temperature (22°C). Animals were sacrificed by asphyxiation with CO_2_ and decapitated at postnatal day 1, 4, 8, 15, 30, 60, 120 and 180. Brains were removed and frozen quickly in liquid nitrogen. Whole brain or brain regions were homogenized in 0.25 M sucrose containing 10 mM Tris pH 8.8 and 1 mM EDTA (STE) using a hand driven pestle. Homogenates were analyzed for specific protein expression (Western blot), mRNA expression (qRT-PCR) and FA analysis (GC-MS) as described below.

### Production of an ACSBG1 knockout mouse (Acsbg1^-/-^)

Acsbg1^-/-^ mice were produced by targeted disruption of exon 2 (encoding amino acid 40-73) of murine Acsbg1. Genomic sequences 5’ and 3’ of Acsbg1 exon 2 were amplified from a mouse genomic bacterial artificial chromosome (BAC) clone containing the Acsbg1 gene, from mouse strain 129vev (Taconic Biosciences). The amplified genomic sequences were sequentially cloned into a vector (ploxpsaβgalneo) containing a β-galactosidase cDNA and a neomycin resistance cassette (Fig. 1A). After confirming the clone orientation by restriction enzyme digestion and DNA sequencing, this plasmid was linearized with Not I and used for transfection of mouse embryonic stem (ES) cells derived from strain 129vev.

**Figure 1.**
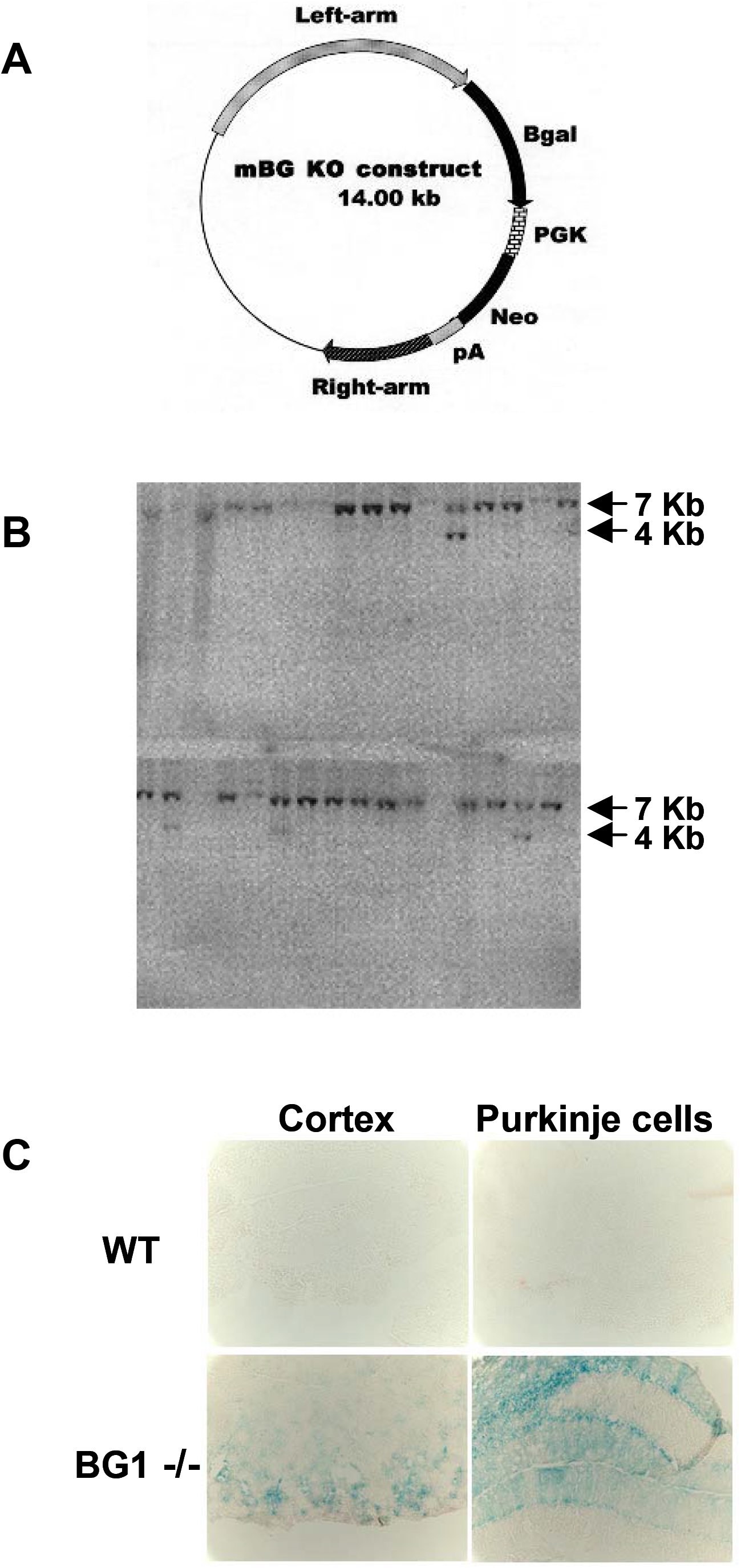
Production of an ACSBG1-deficient (Acsbg1-/-) mouse. A. Knockout (KO) construct. Genomic sequences 5’ and 3’ of Acsbg1 (mBG) exon 2 amplified from a mouse genomic bacterial artificial chromosome were sequentially cloned into a vector containing a β-galactosidase cDNA and a neomycin resistance cassette. B. Southern blot analysis. Mouse embryonic stem (ES) cells were transfected with the linearized KO plasmid. G418-resistant colonies were tested for targeted homologous recombination by Southern blot analysis using a probe for exon 1. The predicted sizes of w.t and KO alleles are 7Kb and 4Kb, respectively. C. β-Galactoside expression. Paraformaldehyde fixed sections of cerebral cortex and cerebellum from w.t. and Acsbg1^-/-^ (BG1-/-) mice were incubated for 2 hrs in X-gal. Cortical neurons and cerebellar Purkinje cells expressing the KO construct stain blue.

Individual colonies resistant to growth in culture medium containing the antibiotic G418 (geneticin) were tested for targeted homologous recombination by Southern blot analysis. Transfected ES cell DNA was digested with BamH1, transferred to a membrane and hybridized with a probe for exon 1. The predicted sizes of the wild type and disrupted Acsbg1 allele are 7Kb and 4Kb, respectively. A representative blot is shown in Fig. 1B. Examination of 150 clones identified 12 with targeted disruption of the Acsbg1 gene. PCR analysis of the insertions confirmed that they were in the proper orientation, i.e., downstream of the Acsbg1 promoter. Cytological analysis revealed 5 clones with normal karyotypes and 4 were used for blastocyst injection. There were 12 chimeras identified on the basis of coat color among the resulting pups. Backcrossing to Blk6 demonstrated that 4 of the chimeras had germline transmission. Acsbg1 heterozygotes were obtained from 2 independent chimeras and Acsbg1^-/-^ homozygotes were generated by brother sister mating. Homozygosity for the disrupted Acsbg1 gene was established by PCR and Southern blot analyses.

### Quantitative real-time PCR assay

Total RNA was extracted from the homogenized cerebellum of WT and Acsbg1^-/-^ mice using the TRIzol® Reagent (Invitrogen, USA) following the manufacturer’s protocol. Total RNA concentration was determined using a NanoDrop 2000 spectrophotometer (Thermo Scientific, USA). Total RNA (3 μg/each reaction) was reverse transcribed into the first-strand cDNA using the SuperScript III First-Strand Synthesis System for RT-PCR kit (Invitrogen, USA) following the manufacturer’s protocol on a Bio-Rad PCR instrument (Bio-Rad, USA). Samples were heated at 65°C for 5 min, 50 °C for 50 min and then 85°C for 5 min. 1 μl of RNase H was added to each tube followed by incubation for 20 min at 37°C. Polymerase chain reaction (PCR) with Taq Polymerase (Invitrogen, USA) was performed using gene-specific primers, designed using Primer 3.0 (Version 0.4.0) software (Table 1). The qRT-PCR reaction with 5 μl cDNA (4ng/μl) and 7.5 μl SYBR®Green PCR Master Mix (5 mL) (Applied Biosystems) was performed on a Bio-Rad CFX connect real-time system (Bio-Rad, USA). After 3 min at 95°C, samples were cycled at 95°C for 10s, 55°C for 10s and 72°C for 30s for 40 cycles; for melt curves sample temperature was increased from 55.0°C to 95.0°C at 0.5°C intervals for 5s each. The GAPDH gene was amplified in the same experiment to serve as the reference gene, and the mRNA expression levels were normalized to those of GAPDH [19].

**Table 1.**
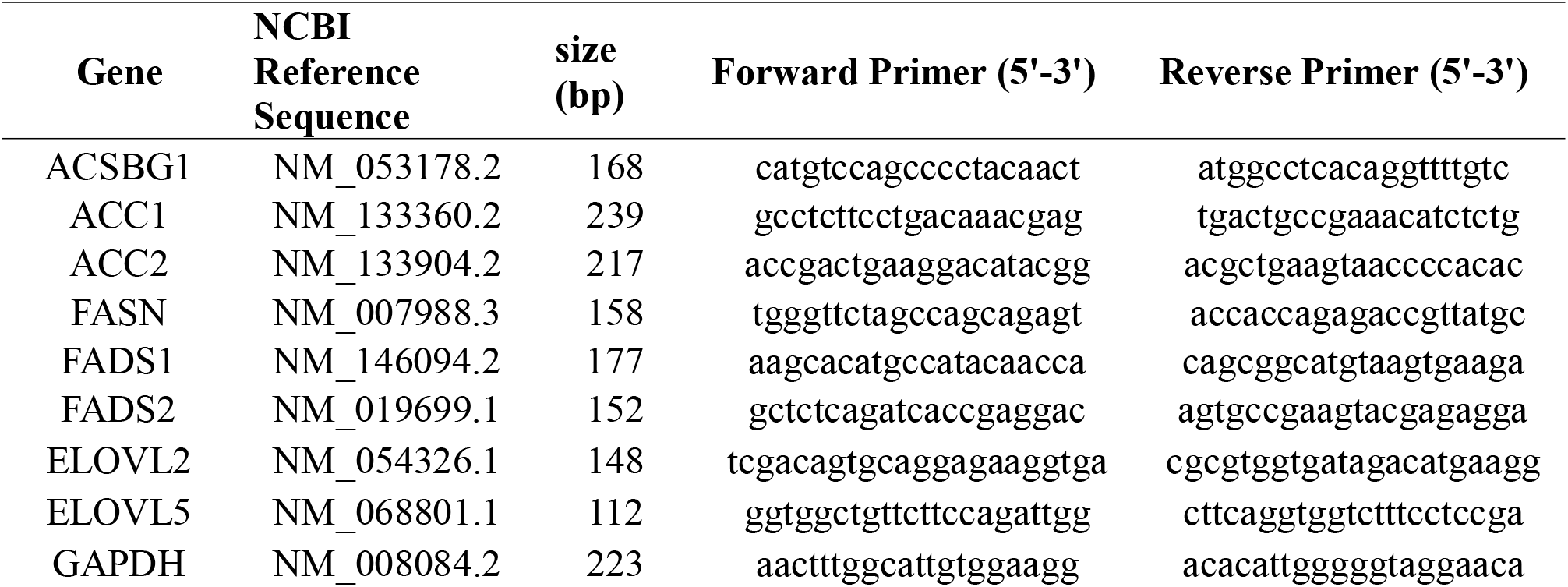
Primer sequences for genes studied.

### Lipid analysis by GC/MS

Fresh cerebellum was harvested from mice (WT and Acsbg1^-/-^) (n=5) of different ages (1, 4, 8, 15, 50, 120 and 180 days) and homogenized in STE using a hand driven pestle. Total fatty acyl groups from 1 mg cerebellar protein were quantitated as their pentafluorobenzyl derivatives on a Capillary Gas Chromatography-Electron-Capture Negative-Ion Mass Spectrometry (GC/MS) system (Agilent, USA) with a Supelco SP2560 capillary column (50m X 0.25mm x 0.2µm) as previously described [20]. Results are expressed as percent of total FA detected in the sample.

### Electrophoresis and Western blotting

Brain or cerebellum from mice of different ages was homogenized in STE (pH8.0) supplemented with protease inhibitors (Roche) and proteins separated by SDS/PAGE. Proteins were transferred to PVDF membranes and analyzed by Western blotting. After blocking with 10% milk for 1 h at room temperature, the membranes were incubated overnight at 4 °C with primary antibody followed by secondary antibody for 1h. Primary antibodies for western blotting were used as follows: anti-rabbit ACSBG1 (1:200); anti-rabbit acetyl-CoA carboxylase (ACC1, 1:4000); anti-rabbit fatty acid synthase (FASN, 1:4000) ); anti-mouse glyceraldehyde-3-phosphate dehydrogenase (GAPDH, 1:20000). Secondary antibodies: goat anti-rabbit horseradish peroxidase (HRP, 1:8000); goat anti-chicken HRP (1:4000); goat anti-mouse HRP (1:8000).

### Statistical analysis

The statistical significance of differences of the biochemical parameters measured in wild-type and Acsbg1^-/-^ mice with different age was determined by Bonferroni’s means comparison test (one way ANOVA), and two sample student’s t –tests. P < 0.05 was considered statistically significant.

## RESULTS

### Production and characterization of an Acsbg1 knockout (Acsbg1^-/-^) mouse

To elucidate the physiological functions of “bubblegum” (ACSBG1), we produced a knockout (KO; Acsbg1^-/-^) mouse by targeted disruption of exon 2 (encoding amino acids 40-73) as described in Methods. Mice were bred to homozygosity, and maintained by mating of Acsbg1^-/-^ males with Acsbg1^-/-^ females. At 21 and 28 weeks of age, Acsbg1^-/-^ mice were ∼18% smaller than w.t. mice of the same age (Table 2). Otherwise, the phenotype of Acsbg1^-/-^ mice was not distinguishable from that of w.t. littermates. Development, behavior, fertility, and lifespan did not appear to be altered by lack of Acsbg1.

**Table 2.** Mouse weight (g ± SD) at 3 and 4 weeks of age.

|  | 21 days | 28 days |
| --- | --- | --- |
| w.t. | 10.0 ± 1.0 | 12.3 ± 1.3 |
| Acsbg1<sup>-/-</sup> | 8.3 ± 0.5 | 10.1 ± 1.1 |

A unique feature of the strategy used to generate this mouse model is replacement of a Acsbg1 exon 2 with the β-galactosidase (*β-GAL*) gene. Thus, tissues that express ACSBG1 in w.t. animals will express *β-GAL* in KO mice, and thus be readily identified. Therefore, we assessed expression of the *β-GAL* gene in the brain of an Acsbg1^-/-^ mouse. Sections of brain were fixed in 4% paraformaldehyde and incubated for 2 hrs in X-gal (5-bromo-4-chloro-3-indolyl-β-D-galactosidase) prior to microscopic examination. As seen in Fig. 1C, β-galactosidase activity could be detected in cerebral cortical neurons and cerebellar Purkinje cells, known sites of ACSBG1 expression [13]. This result is consistent with β-galactosidase expression being driven by the Acsbg1 gene promoter.

Previous studies used indirect immunofluorescence (IF) to show that, in addition to cortical neurons and Purkinje cells, ACSBG1 was expressed in cortisol-producing cells of adrenal gland, testosterone-producing cells of testis and estrogen-producing cells [13,21]. Lack of ACSBG1 expression in these tissues in the Acsbg1^-/-^ mouse was confirmed by Western blotting. ACSBG1 protein was detected in adrenal gland, testis, and brain from w.t. mice, but was not seen in Acsbg1^-/-^ mice (Fig 2. No expression of ACSBG1 was detected in liver, heart, kidney, lung, or spleen from either w.t. or Acsbg1^-/-^, as expected (Fig. 2).

**Figure 2.**
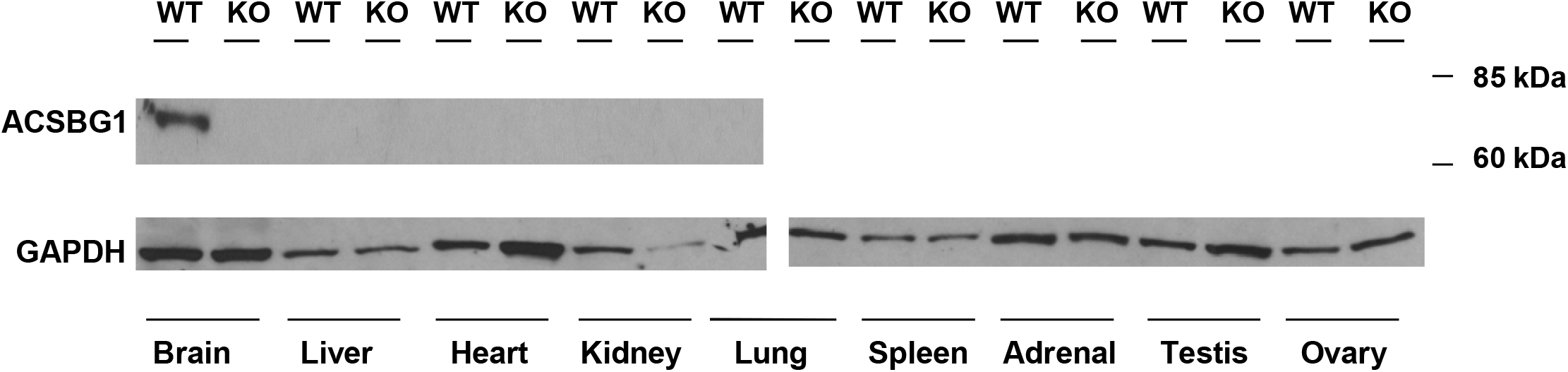
Expression of ACSBG1 in mouse tissues. Tissues from adult w.t. and Acsbg1^-/-^ (KO) mice were homogenized and subjected to Western blot analysis as described in Methods. The ∼70 kDa ACSBG1 band was observed in brain, adrenal gland, testis and ovary of w.t. mice. GAPDH was used as a loading control.

### Acyl-CoA synthetase activity in w.t. and Acsbg1^-/-^ mouse tissues

Lack of a specific acyl-CoA synthetase might be expected to lower enzyme activity towards FA known to be substrates in tissues normally expressing the ACS. We previously demonstrated that endogenous ACSBG1 preferred the long-chain FA palmitic acid (C16:0), but only weakly activated the very long-chain FA lignoceric acid (C24:0) [12,14]. Therefore, we prepared tissue homogenates and measured their ability to activate these FA. Because endogenous ACSBG1 in Neuro2a cells sedimented with a mitochondria-enriched fraction [13], we also prepared and assayed ACS activity in brain mitochondria. As shown in Table 3, lack of ACSBG1 failed to lower C16:0 activation in whole brain, cerebellum, brain mitochondria, adrenal gland, or testis. In fact, C16:0 activation in cerebellum and adrenal glands of Acsbg1^-/-^ mice was higher than that measured in w.t. mice. As expected, there was no significant change in ACS activity toward the C24:0 substrate when ACSBG1 was depleted. Similarly, ACS activity with either substate was not changed by depletion of ACSBG1 in liver (data not shown).

**Table 3.** Tissue ACS activity of w.t. and Acsbg^-/-^ mice. Tissues were collected and homogenized in STE. The ability to activate long- or very long-chain FA to their CoA derivatives was measured as described in Methods using radiolabeled palmitic acid (C16:0) or lignoceric acid (C24:0), respectively, as substrate.

| Tissue | C16:0 activation<br>(nmol/20 min/mg protein) |  | C24:0 activation<br>(nmol/20 min/mg protein) |  |
| --- | --- | --- | --- | --- |
|  | w.t. | Acsbg1<sup>-/-</sup> | w.t. | Acsbg1<sup>-/-</sup> |
| Whole brain | 16.5 | 17.3 | 2.2 | 2.0 |
| Cerebellum | 8.7 | 16.5 | 0.6 | 0.8 |
| Brain mitochondria | 26.9 | 26.7 | 5.5 | 5.3 |
| Adrenal gland | 38.2 | 50.0 | 1.2 | 1.4 |
| Testis | 18.8 | 18.6 | 1.7 | 1.6 |

### Regional distribution of ACSBG1 expression in mouse brain

To determine semi-quantitatively which brain regions normally express ACSBG1, brains from Acsbg1^-/-^ and w.t. mice were collected and dissected into cortex, hippocampus, white matter, brainstem and cerebellum. Western blotting revealed that ACSBG1 (∼70 kDa band) was expressed in all brain regions in w.t. mice (Fig. 3A). Expression was significantly higher (p<0.01) in cerebellum than in the other four regions (Fig. 3B); cerebellar expression was nearly 5-fold higher than that in cortex. No ACSBG1 protein expression was detectable in any of the Acsbg1^-/-^ mouse brain regions examined.

**Figure 3.**
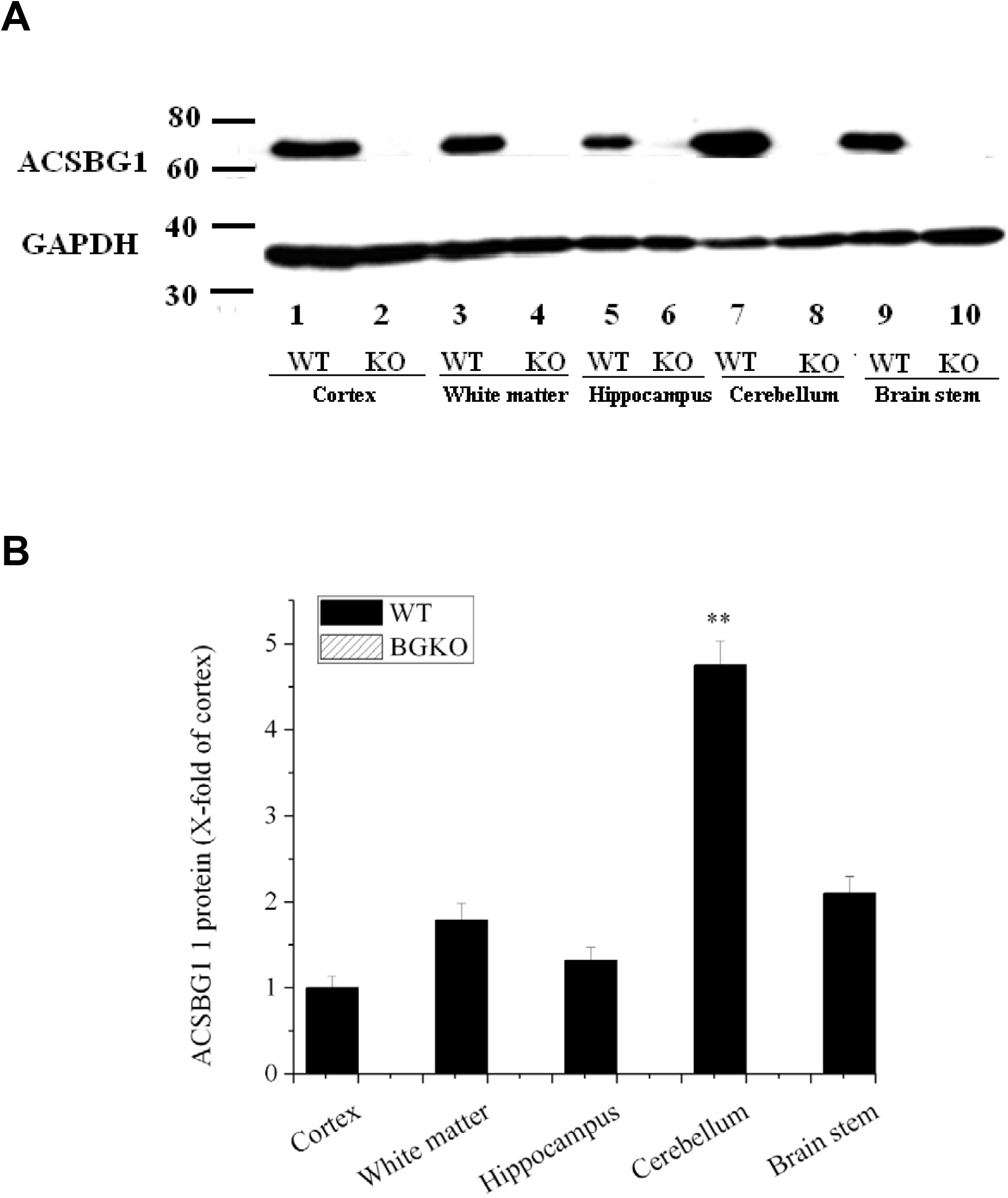
Regional expression of ACSBG1 in mouse brain. A. Western blotting. Brain from adult w.t. and Acsbg1^-/-^ (KO) was dissected into cortex, hippocampus, white matter, brainstem and cerebellum. Western blotting revealed that ACSBG1 (∼70 kDa band) was expressed in all brain regions in w.t. mice. No ACSBG1 expression was detected in KO mice. GAPDH was used as a loading control. B. Relative expression of ACSBG1 in brain regions. Densitometry was used to quantify Western blot data. Results were normalized to GAPDH expression and are presented as fold-increase relative to cortex.

### Brain ACSBG1 expression increases with development of w.t. mice

The average age of onset of neurological symptoms in CCER is around 6 years of age, whereas symptoms of AMN typically begin in early adulthood [4]. We therefore wanted to determine the developmental expression pattern of ACSBG1 in mouse brain. Since cerebellum had the highest ACSBG1 expression among the regions examined, we used quantitative PCR and Western blotting to detect Acsbg1 mRNA and protein levels, respectively, in this area of the brain. In the cerebellum of mice, Acsbg1 mRNA increased with age, and reached maximum in 1-2-month-old mice (Fig 4A). For ACSBG1 protein expression, there was a very low expression level in 1, 4 and 8 day-old mice, which increased dramatically with further development and peaked at around 2 months (Fig 4B, p<0.01 vs. 1 day-old). At this age, the Acsbg1 mRNA level was about 5 times higher than that found in 1-day-old mice, and the amount of protein expressed was about 100-fold higher. As the mice aged, Acsbg1 mRNA and protein expression in cerebellum decreased slightly. No Acsbg1 mRNA or protein was detected in cerebella from Acsbg1^-/-^ mice at any time point, further confirming total absence of ACSBG1 in this mouse model.

**Figure 4.**
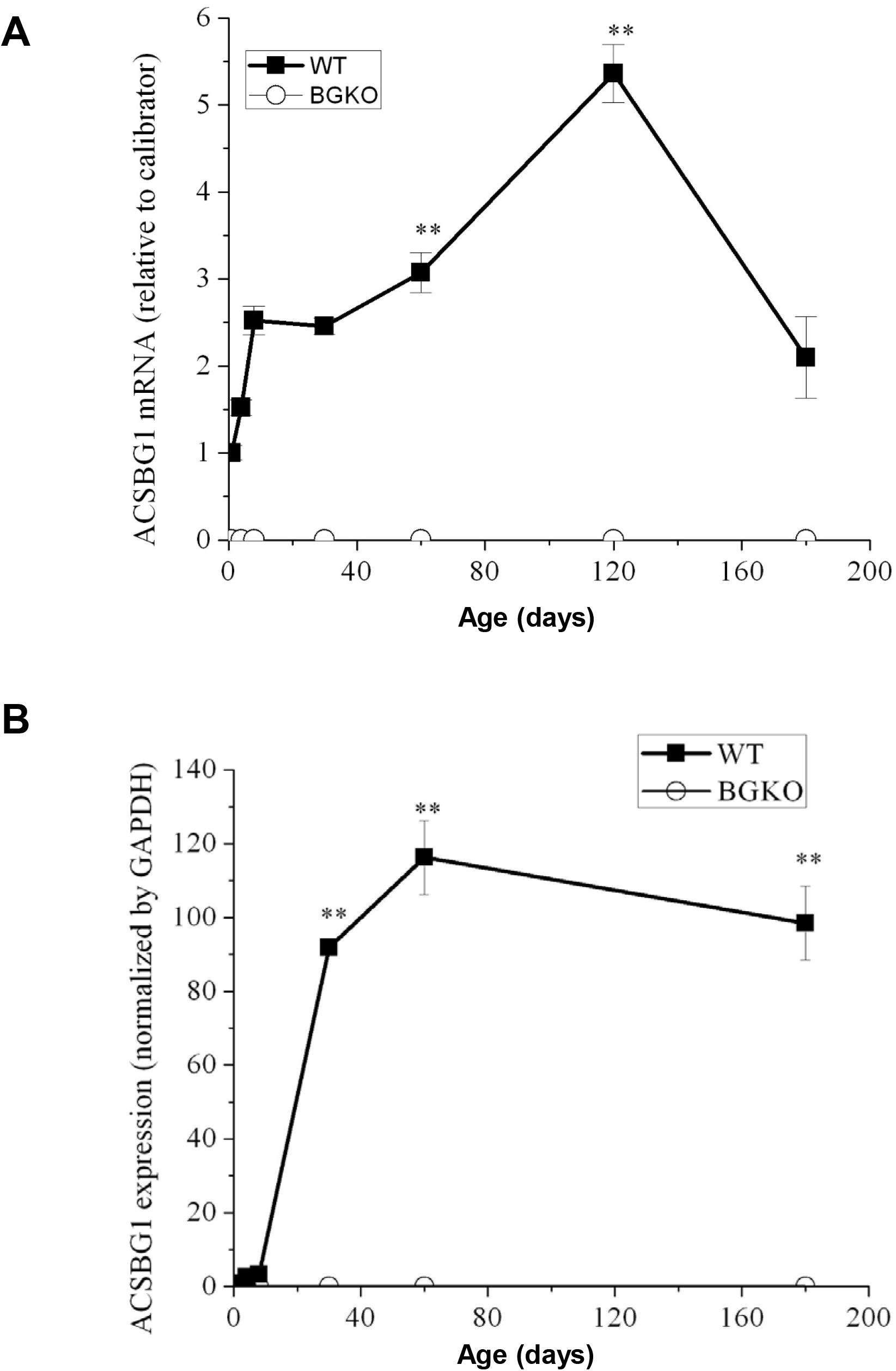
Developmental expression of ACSBG1 in cerebellum. A. mRNA expression. Acsbg1 mRNA in cerebellum from w.t. and Acsbg1^-/-^ mice of increasing age was measured by quantitative PCR as described in Methods. No Acsbg1 mRNA was detected in KO mouse cerebellum. B. Western blotting. ACSBG1 protein expression in cerebellum of mice of increasing age was quantitated by densitometric analysis of the Western blot shown in Supplemental Figure 2. Results were normalized to GAPDH expression. No ACSBG1 protein was detected in KO mouse cerebellum.

### ACSBG1 depletion decreases levels of saturated VLCFA and monounsaturated FA while increasing ω3 polyunsaturated FA levels in cerebella of adult mice

The defining biochemical abnormality in XALD is elevated levels of saturated VLCFA [4]. To assess whether lack of ACSBG1 affected brain FA levels and composition during development, cerebella from w.t. and Acsbg1^-/-^ mice of different ages were homogenized, their lipids extracted, and the FA composition measured by GC/MS as described in Methods. We first looked at differences between classes of fatty acids, e.g. saturated, ω9 monounsaturated, ω6 polyunsaturated, in w.t. and Acsbg1^-/-^. As shown in Table 4 and Suppl. Fig. 3, total saturated FA (C10-30, which constitute nearly half of total FAs) were mainly unchanged with age but tended to be a bit lower in Acsbg1^-/-^ mice. When we looked only at VLCFAs (C22-30; ∼3% of total FAs in adult mice), we found that levels of these FA were barely detectable at birth and increased relatively linearly over the first 50 days of life. Levels in Acsbg1^-/-^ cerebella were lower than in w.t. mice. From 50-180 days, saturated VLCFA slightly increased in w.t. mice while remaining the same or decreasing somewhat in Acsbg1^-/-^ mice. The lower VLCFA in ACSBG1-deficient mice is thus in contrast to the increased VLCFA seen in XALD mice.

**Table 4.** Change in cerebellar FA levels in w.t. and Acsbg^-/-^ mice with age. Cerebellum was collected from w.t. and Acsbg1^-/-^ mice at 1, 4, 8, 15, 50, 120, and 180 days of age. Lipids were extracted and total lipid FAs were quantitated as described in Methods. Levels of FA in different classes are shown as a percentage of total FA. These data are also shown graphically in Suppl. Fig. 3.

| Age (days) |  | 1 | 4 | 8 | 15 | 50 | 120 | 180 |
| --- | --- | --- | --- | --- | --- | --- | --- | --- |
| Total saturated FA | w.t. | 47.5% | 48.6% | 47.0% | 47.3% | 46.4% | 45.9% | 44.9% |
|  | Acsbg1<sup>-/-</sup> | 40.4% | 47.2% | 47.1% | 47.0% | 45.3% | 45.7% | 43.4% |
| Total sat VLCFA (C22-30) | w.t. | 0.2% | 0.2% | 0.3% | 1.2% | 2.7% | 2.8% | 3.2% |
|  | Acsbg1<sup>-/-</sup> | 0.1% | 0.2% | 0.4% | 1.0% | 2.5% | 2.3% | 2.4% |
| Total ω9 FA | w.t. | 14.9% | 13.8% | 14.2% | 15.9% | 21.3% | 21.7% | 24.2% |
|  | Acsbg1<sup>-/-</sup> | 15.3% | 13.6% | 13.8% | 14.6% | 19.0% | 20.3% | 21.0% |
| Total ω5 +ω7 FA | w.t. | 5.4% | 4.9% | 5.2% | 4.3% | 4.3% | 4.1% | 4.4% |
|  | Acsbg1<sup>-/-</sup> | 5.2% | 5.3% | 4.9% | 4.4% | 4.6% | 4.2% | 4.6% |
| Total ω6 FA | w.t. | 17.8% | 18.5% | 18.7% | 18.0% | 12.5% | 12.4% | 11.8% |
|  | Acsbg1<sup>-/-</sup> | 21.2% | 18.0% | 19.0% | 18.8% | 13.5% | 13.0% | 12.8% |
| Total ω3 FA | w.t. | 11.6% | 11.7% | 12.7% | 13.6% | 15.3% | 15.7% | 14.7% |
|  | Acsbg1<sup>-/-</sup> | 15.0% | 13.4% | 13.2% | 14.0% | 17.6% | 16.7% | 18.1% |
| Total trans FA | w.t. | 2.9% | 2.6% | 2.2% | 0.9% | 0.2% | 0.2% | 0.2% |
|  | Acsbg1<sup>-/-</sup> | 2.9% | 2.7% | 2.1% | 1.2% | 0.1% | 0.2% | 0.1% |

In both w.t. and Acsbg1^-/-^ mice, total ω9 monounsaturated FA (constituting ∼20% of total FAs in adult mice) increased slightly over the first 50 days of life and then remained relatively constant through 180 days. At every time point, levels were somewhat higher in w.t. than in Acsbg1^-/-^ cerebella. Levels of ω5+ω7 monounsaturated FAs (∼5% of total FAs) decreased over the first fifteen days of life and then remained relatively constant; at all ages levels were similar in both groups of mice. The ω6 polyunsaturated FAs (∼12% of total FAs in adults) decreased over the first 50 days of life before remaining constant thereafter; levels of these FA were consistently slightly higher in ACSBG1-deficient mice. Interestingly, levels of ω3 polyunsaturated FAs (∼15% of total FA in adults) were also consistently higher in knockout mice relative to w.t. animals; levels rose slightly over the first 50 days of life and did not change appreciably thereafter. Total trans-FAs decreased from about 3% of total FAs to nearly zero over the first 50 days and were barely detectable thereafter; levels in w.t. and ACSBG1-deficient cerebella were not different.

### Levels of specific FA are affected by ACSBG1 depletion in cerebella of adult mice

Not all FA of a specific class, e.g. monounsaturated FA, were affected by lack of ACSBG1. In Figure 5, age-related changes in the levels of several FA are shown. Some are typical members of a given class, while others are specific members that show changes in Acsbg1^-/-^ mice. Levels of the two most abundant saturated FA in nature – C16:0 (palmitic acid) and C18:0 (stearic acid) – were unchanged by ACSBG1 knockout. In contrast, levels of saturated very long-chain FA containing 24 (lignoceric acid) and 26 carbons (cerotic acid) were significantly lower in Acsbg1^-/-^ mice. Among the ω9 monounsaturated FA, levels of both the abundant C18:1 (oleic acid) and the very long-chain C24:1 (nervonic acid) were lower in cerebella of Acsbg1^-/-^ mice. Interestingly, both saturated and monounsaturated VLCFA increased from nearly zero at birth to steady state levels at around 2 months of age. In contrast, levels of both C16:1ω9 and C16:1ω7 monounsaturates decreased from more than 2% of total FA at birth to very low (ω7) and nearly undetectable (ω9) levels at 2 months of age; there were no differences between w.t. and Acsbg1^-/-^ mice.

**Figure 5.**
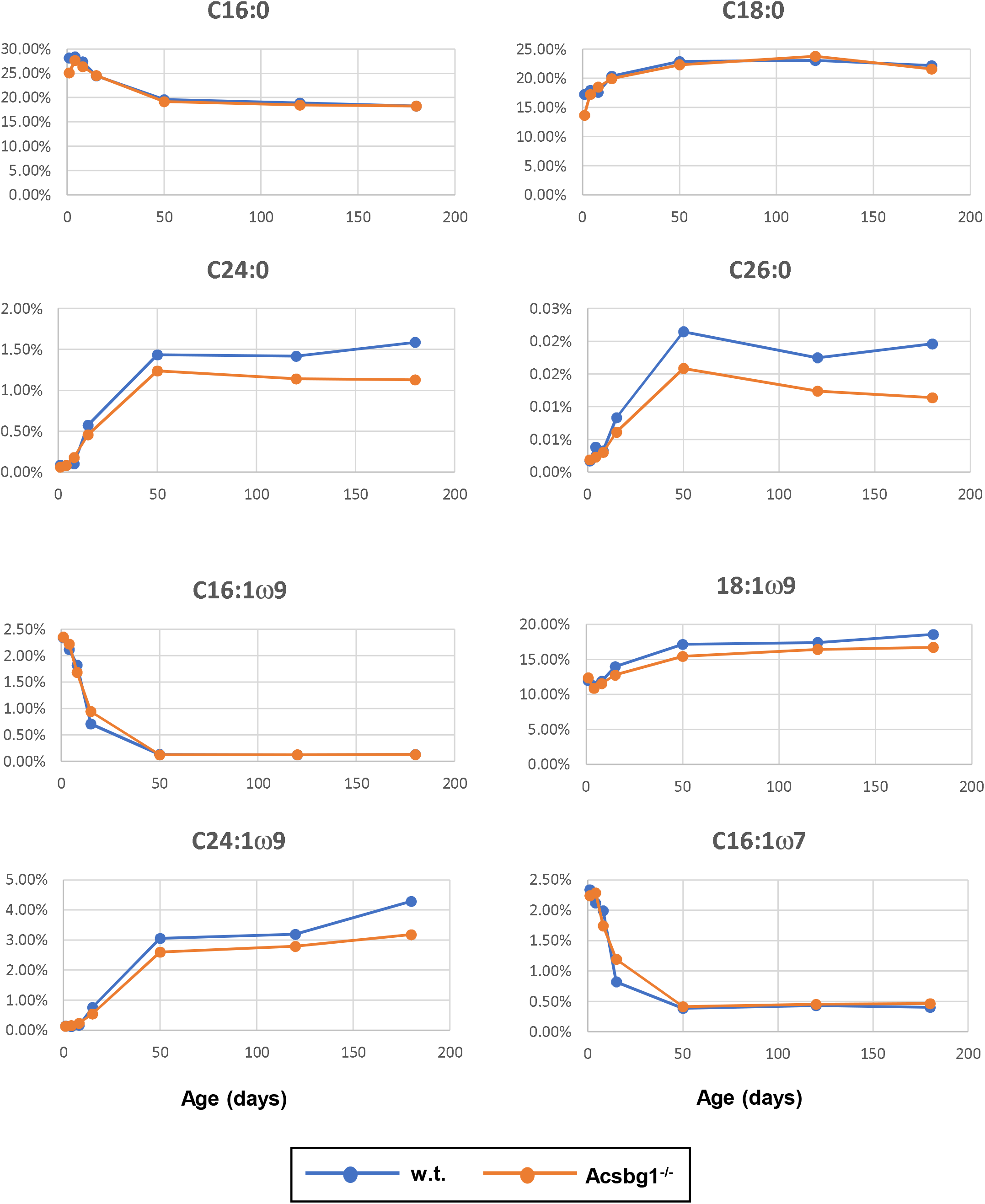
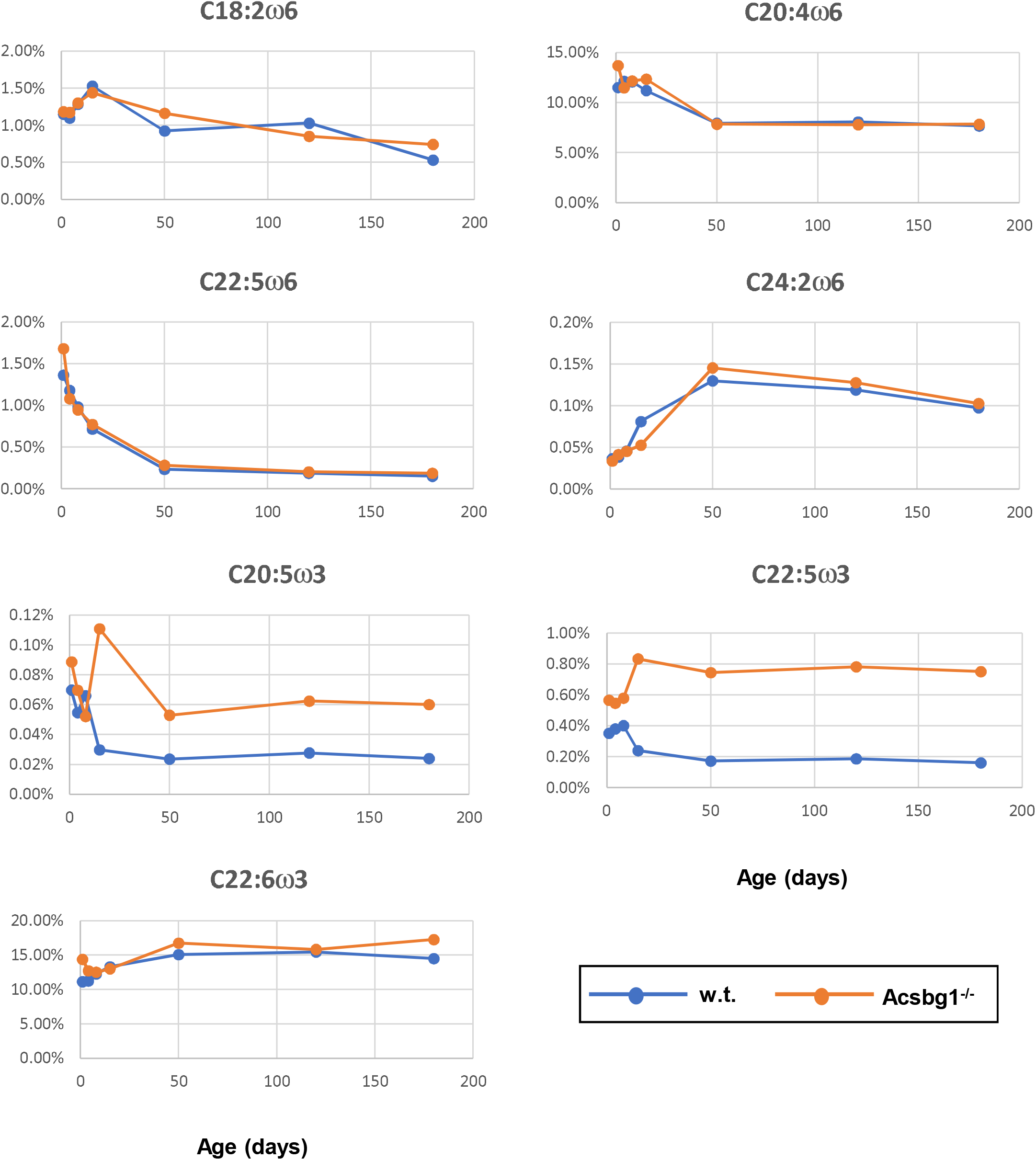
Change with age in levels of specific cerebellar FA in w.t. and Acsbg-/- mice. Lipids were extracted from cerebellum and levels of specific FA quantitated as described in the legend to Table 4. Levels of specific FA as a percentage of total FA are shown.

Among the ω6 polyunsaturated FA, none showed a significant difference in level between w.t. and Acsbg1^-/-^ mice. There were, however, some age-related changes. Plots of FA level vs. age are shown for C18:2 (linoleic acid), C20:4 (arachidonic acid), C22:5 (docosapentaenoic acid), and C24:2.

Lack of ACSBG1 did produce changes in levels of some, but not all, polyunsaturated FA of the ω3 series. Specifically, levels of C20:5 (eicosapentaenoic acid) and C22:5 were significantly higher in Acsbg1^-/-^ mice.

### Depletion of ACSBG1 affects the developmental expression pattern of enzymes required for de novo FA synthesis in mouse cerebellum

Differences in the FA profile of cerebella from w.t. and Acsbg1-/- mice (Table 4, Figure 5, and Suppl. Figure 3) suggest that the presence or absence of ACSBG1 has effects on lipid metabolism during mouse brain development. To understand better the role of ACSBG1 in these processes in cerebellum, we quantified several enzymes relevant to FA synthesis and degradation by qRT-PCR and western blotting in w.t. and Acsbg1^-/-^ mice. Acetyl-CoA carboxylase 1 (ACC1) catalyzes the regulated initial step in *de novo* FA synthesis – carboxylation of acetyl-CoA to form malonyl-CoA [22]. As shown in Figure 6A, ACC1 mRNA in cerebella of w.t. mice decreased over the first 6 months of life, and then slowly increased over the next 5 months. In ACSBG1-deficient mice, ACC1 mRNA was significantly higher than in w.t. mice in the early postnatal period, but by 2-3 weeks of age levels were similar.

**Figure 6.**
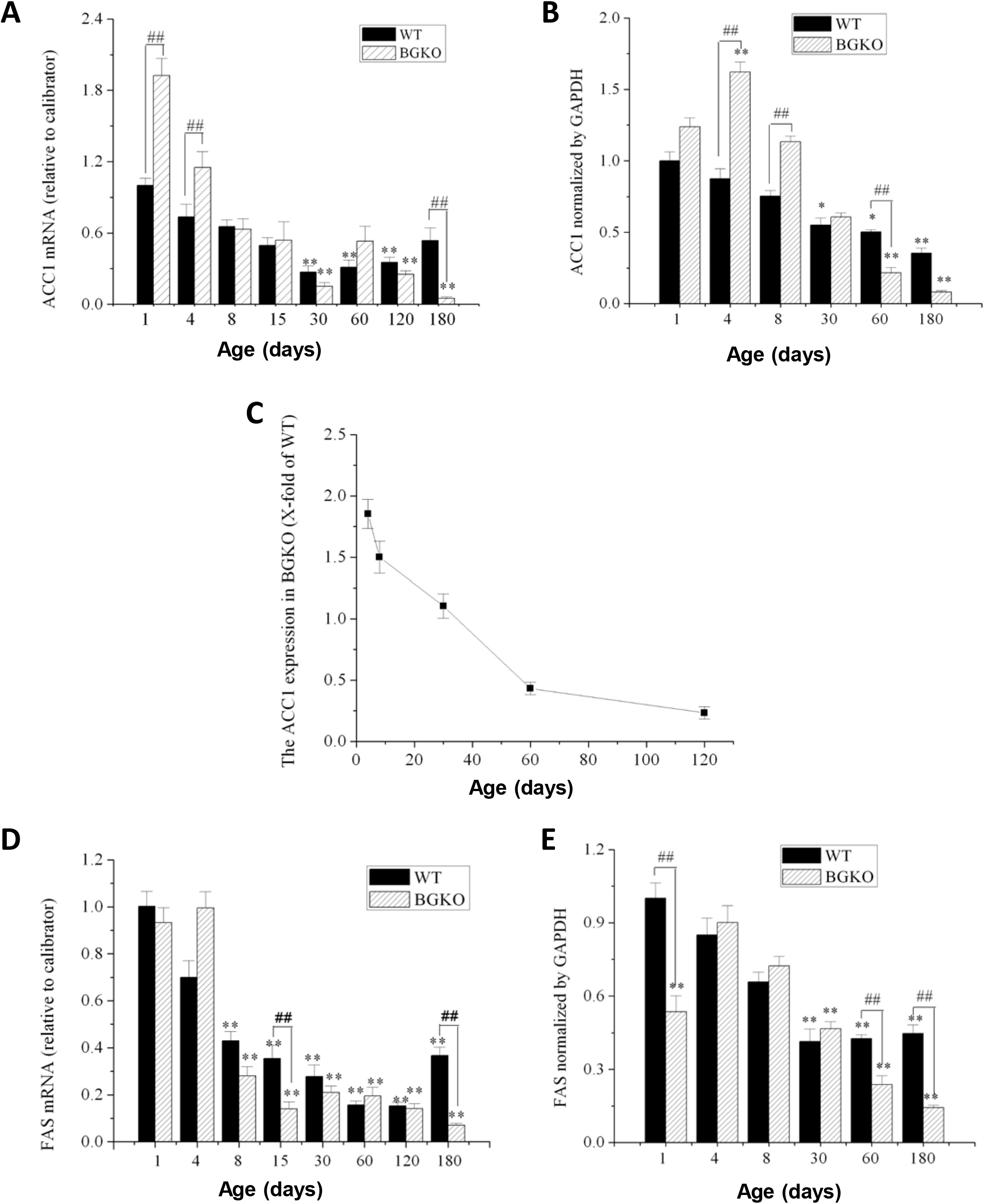
Developmental expression of de novo FA synthesis enzymes in cerebellum from w.t. and Acsbg-/- mice. Quantitative PCR was used to measure level of ACC1 (A) and FASN (D) mRNA in cerebellum obtained from mice at the indicated age. Expression relative to that in 1 day-old w.t. mice is shown. Western blotting was used to quantitate protein expression of ACC1 (B) and FASN (E) in cerebellum of w.t and ACSBG1 KO mice. Densitometry of ∼70 kDa ACSBG1 bands on blots was normalized to GAPDH. Protein levels relative to that seen in 1 day-old w.t. mice are shown. To analyze further the developmental progression of ACC1 protein expression, a derivative plot of the data shown in panel B was created by dividing KO mouse expression by w.t. expression. This plot is shown in panel C.

However, while ACC1 expression slowly rebounded in w.t. mice, mRNA levels continued to decline in Acsbg1^-/-^ mice so that by 3-6 months of age, mRNA levels were significantly lower in Acsbg1^-/-^ mice than in w.t. mice. In addition to mRNA, we also measured ACC1 protein expression by densitometric scanning of Western blots. In general, ACC1 protein quantitation in w.t. and Acsbg1-/- mice paralleled Acc1 mRNA levels (Figure 6B). The time-dependent decrease in ACC1 protein level was more linear than what was seen with mRNA levels, and the drop in ACSBG1-deficient mouse cerebellum was more acute than in w.t. (Figure 6C).

Fatty acid synthase (FASN) is a large multienzyme complex whose component domains catalyze all reactions of *de novo* FA synthesis downstream of ACC1 [23]. Like ACC1, cerebellar Fasn mRNA was high in the early postnatal period of w.t. mice and decreased to a low, steady state by 1-2 months of age (Figure 6D). In Acsbg1^-/-^ mice, mRNA levels also started high but decreased a bit more rapidly than in w.t. mice; at 6 months of age Fasn mRNA levels were lower in Acsbg1^-/-^ mice. FASN protein expression in general paralleled mRNA expression in both w.t. and Acsbg1^-/-^ mice (Figure 6E).

### mRNA expression of several other FA metabolism enzymes is affected by ACSBG1 deficiency

We measured mRNA expression of several additional enzymes involved in FA metabolism in cerebella from w.t. and Acsbg1^-/-^ mice. While ACC1 is essential for de novo FA synthesis in lipogenic tissues, its isoform ACC2 produces malonyl-CoA primarily to regulate FA degradation by inhibition of carnitine palmitoyltransferase 1 (CPT1) and carnitine octanoyltransferase (CROT) in mitochondria and peroxisomes, respectively [22,24]. Acc2 mRNA levels were quantitated by RT-PCR in cerebellum from w.t. and Acsbg1^-/-^ mice (Figure 7A). In general, there was no major change during mouse development and no significant difference between w.t. and Acsbg1^-/-^ mice.

**Figure 7.**
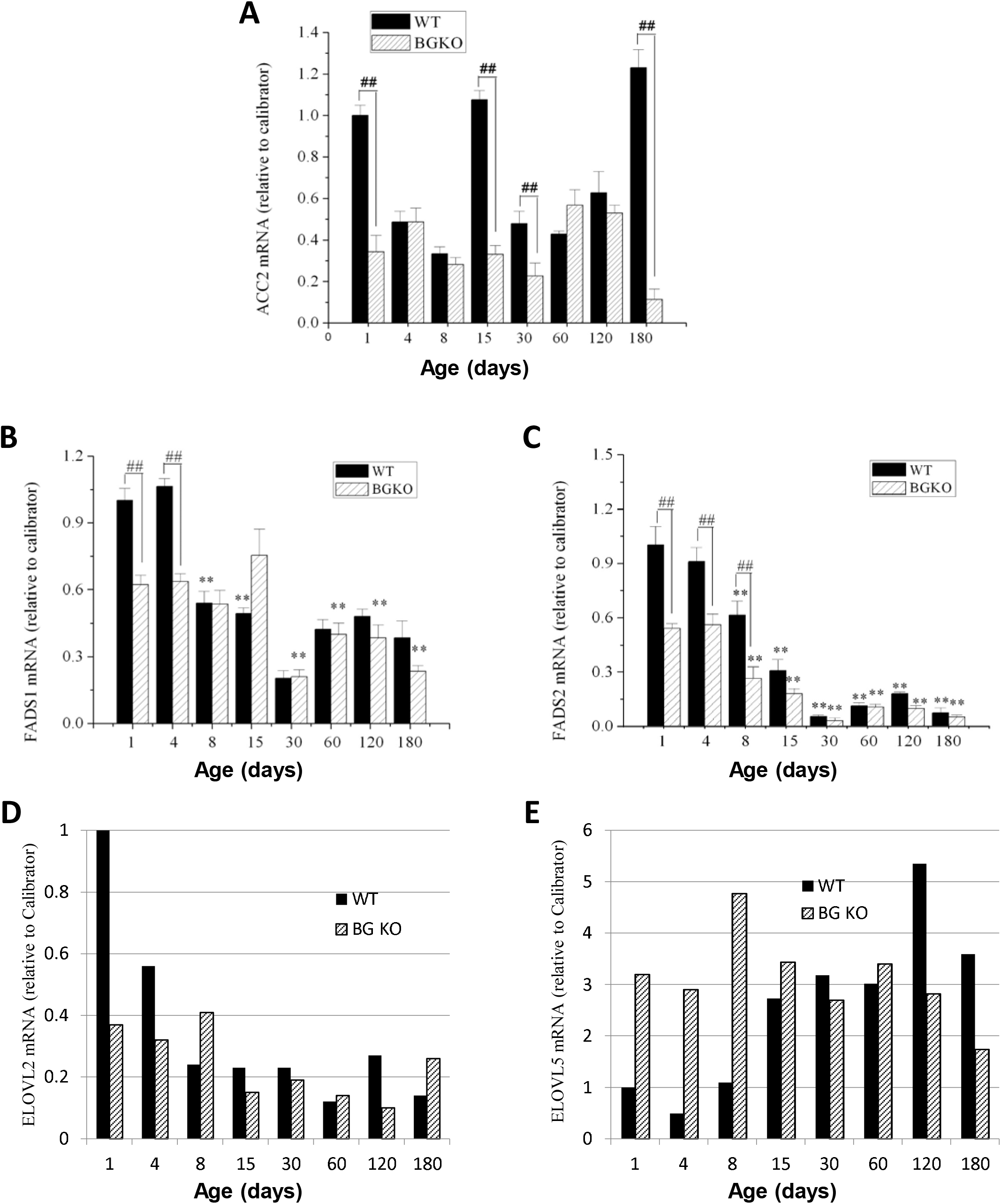
Developmental expression of enzymes participating in FA degradation, elongation and unsaturation in cerebellum from w.t. and Acsbg-/- mice. Quantitative PCR as described in the legend to Fig. 6 was used to measure mRNA levels of several additional FA metabolism enzymes, including: (A) ACC2, a regulator of fatty acid entry into mitochondria for degradation by β-oxidation, (B) FADS1 and FADS2, desaturases relevant to synthesis of ω3 polyunsaturated FA; (D) ELOV2 and (E) ELOVL5, elongases that are also relevant to ω3 polyunsaturated FA synthesis.

FA desaturases FADS1 and FADS2 participate in the synthesis of polyunsaturated FA by insertion of double bonds 5 or 6 carbons (respectively) from the carboxyl carbon [25]. The mRNA levels of both desaturases decreased with increasing age of w.t. mice, reaching a relative steady state by around one month of age (Figure 7B and 7C). The steady state level of Fads2 mRNA, however, was significantly lower than that seen with Fads1 mRNA.

FA elongases (ELOVL family) catalyze the initial step in adding 2-carbon units to long chain FA to produce very long-chain FA [26]. In particular, ELOVL2 and ELOVL5 work in concert with the desaturases to produce polyunsaturated FA of increasing chain lengths [26]. Elovl2 mRNA levels in w.t. mice were high at birth but rapidly decreased to a rather steady state level by 2 weeks of age (Figure 7D). In Acsbg1^-/-^ mice, mRNA levels were lower than w.t. at birth and did not change appreciably throughout life. On the other hand, levels of Elovl5 mRNA in cerebellum from w.t. mice were low at birth, but increased and reached a relative steady state by about 2 weeks of age (Figure 7E). In Acsbg1^-/-^ mice, Elovl5 mRNA levels were significantly higher at birth and remained rather constant throughout life. However, in 4-6 month old mice, w.t. mRNA levels were higher in w.t. than in Acsbg1^-/-^ mice.

## DISCUSSION

Amongst the inherited metabolic diseases, XALD is one of the most prevalent, with an allele frequency of about 1 in 17,000 [27]. Nearly all male patients have some degree of adrenal gland insufficiency. While some will develop central demyelination in childhood, the CCER XALD phenotype, most will manifest symptoms of peripheral nervous system disease, adrenomyeloneuropathy (AMN) in early adulthood [4]. Progression to the CCER phenotype in later adulthood is seen in more than 50% of AMN patients [28]. In addition, some AMN patients show signs of testicular involvement including decreased spermatogenesis or infertility [4]. Thus the pathology in XALD is limited to a relatively small number of tissues. In contrast, ABCD1, the gene defective in XALD, has a rather ubiquitous tissue expression profile (Suppl. Figure 1). ABCD1 expression is highest in adipose tissue and small intestine - tissues that have not been shown to be clinically affected in XALD. While testis and adrenal gland have the next highest ABCD1 expression, levels of brain ABCD1 are among the lowest found in human tissues. Why, therefore, is pathology in XALD limited to only a few tissues?

A definitive explanation for this remains elusive. One hypothesis is that the involved tissues have unique metabolic features that render them more vulnerable to the high levels of VLCFA that are the biochemical signature of XALD [29,30]. Several properties of ACSBG1, identified in a screen of fruit fly neurodegeneration mutants [3], suggested that exploration of its potential role in XALD pathogenesis was warranted. When first discovered, ACSBG1 was a previously unknown member of the ACS enzyme family. ACSs occupy a central position in lipid metabolism by activating FA to their CoA derivatives, a necessary prerequisite for subsequent participation in either catabolic or anabolic processes [1]. In addition to neurodegeneration, ACSBG1-deficient flies had elevated tissue levels of VLCFA, similar to the situation in XALD patients [4]. We and others found that in humans and mice, expression of ACSBG1 was primarily in the tissues pathologically affected in XALD, namely brain, adrenals, and testis [13,14]. These observations prompted us to create a knockout mouse model to test the relationship between ACSBG1 and XALD.

The Acsbg1^-/-^ mouse described herein was generated by replacement of exon 2 with a *β-GAL* gene. Sheng et al. [31] created a similar knockout mouse by deletion of a ∼2 kb fragment of the Acsbg1 gene that included the start codon; these researchers originally called this gene “gonadotropin-regulated long chain fatty acid Acyl-CoA synthetase” (GR-LACS). No gross phenotypic abnormalities were observed in either our Acsbg1^-/-^ mouse or the GR-LACS^-/-^ mouse. We did observe a mild growth phenotype that was not present in the GR-LACS^-/-^ mouse. Lack of ACSBG1 did not reduce ACS activity with either C16:0 or C24:0 as substrate in tissues from Acsbg1^-/-^ mice produced by either laboratory, suggesting that there was compensatory upregulation of other ACS genes. Studies of Acsbg1^-/-^ mice from both laboratories confirmed that tissues expressing ACSBG1 were identical to those pathologically affected in XALD.

When we looked at the relative expression of ACSBG1 in various brain regions, we observed that the highest expression was in cerebellum (Figure 3). To probe further the potential relevance of ACSBG1 in XALD, we looked at expression of this protein during brain developmental. We found ACSBG1 to be very low at birth and through the first few days of life, after which there was a robust increase in expression (Figure 4). In contrast, expression of ABCD1 was reported to be at its highest from embryonic day 12 through postnatal day 15, after which expression dropped 2.6-fold by adulthood [32]. Thus, there was little correlation between expression of the two proteins with age.

Because Drosophila *bubblegum* mutants had elevated levels of saturated VLCFA [3], similar to XALD patients [4], we expected that Acsbg1^-/-^ mice would also exhibit a similar elevation, particularly in brain. While cerebellar total saturated FA levels in w.t. and Acsbg1^-/-^ mice were essentially the same from birth to 6 months of age, VLCFA levels differed. However, instead of the predicted increase in VLCFA levels in Acsbg1^-/-^ mouse cerebellum, saturated VLCFA levels were consistently lower (Table 4 and Suppl. Fig. 3). This observation decreases the likelihood that ACSBG1 has a direct role in XALD pathophysiology. Interestingly, a pattern similar to cerebellar VLCFA levels in w.t. vs. Acsbg1^-/-^ mice was seen for total ω9 monounsaturated FA, and the reverse was detected for total polyunsaturated ω3 FA where levels in Acsbg1^-/-^ mice were higher than in w.t.

When we looked at individual cerebellar FA levels in w.t. and Acsbg1^-/-^ mice from birth through 6 months of age (Fig. 5), a few comparisons were notable. Since C16:0 (palmitic acid) is a preferred substrate for ACSBG1 [12,14], it was predicted that levels of this FA would be lower in tissues from Acsbg1^-/-^ mice. However, essentially no differences between w.t. and Acsbg1^-/-^ mouse C16:0 content were noted throughout the first 6 months of life. A similar pattern was seen with C18:0 (stearic acid). In contrast, by two months of age, saturated VLCFAs C24:0 and C26:0 were clearly lower in Acsbg1^-/-^ mouse cerebellum, and this gap grew wider with age to at least 6 months. A similar pattern was seen with the ω9 monounsaturated VLCFA, C24:1 (nervonic acid), and to a lesser extent with C18:1ω9 (oleic acid).

The increase in total ω3 polyunsaturated FAs noted above was primarily due to increases in C22:6ω3 (docosahexaenoic acid; DHA). Brain and cerebellar DHA levels are surprisingly high, accounting for up to 25% of total brain FA in humans [33]. Although C20:5ω3 (eicosapentaenoic acid; EPA) and C22:5ω3 (docosapentaenoic acid; DPA) are intermediates in the synthesis of DHA [34], it seems unlikely that the increased EPA and DPA contribute significantly here, as the levels of these FA are about 3- and 2-orders of magnitude lower (respectively) than DHA. A large number of previous studies indicated that ω3 polyunsaturated FA are essential for normal growth and development. The health effects of these FAs include reduction of cardiovascular risk due to antiarrhythmic, anti-inflammatory, anti-thrombotic and lipid lowering actions, as well as improved glucose level control and insulin sensitivity, and neuroprotection [34–37]. Thus, reduced ACSBG1 expression could be associated with improvement in cardiovascular disease, reduced complications of diabetes and lowered risk for depression. These effects could be mediated by eicosanoids and/or docosanoids derived from these ω3 polyunsaturated FAs, where even very low levels of signaling molecules can have highly significant biological activity. Further investigation into the effect of ACSBG1 depletion on eicosanoid- and docosanoid-mediated signaling in brain is clearly warranted.

Sheng et al. also measured FA levels in adult brain, testis, ovary and adrenal in their Acsbg1^-/-^ mouse [31]. Our findings in cerebellum and their findings in whole brain are in general agreement for many FA, including C16:0, C18:1, C24:1, C20:5ω3, C22:5ω3 and C22:6ω3.

The long-chain saturated FA palmitate (C16:0) is produced by the *de novo* FA synthesis pathway that includes acetyl-CoA carboxylase (ACC1) and fatty acid synthase (FASN) [22,23]. C16:0 can then be elongated to produce VLCFA. At and shortly after birth, cerebella from Acsbg1^-/-^ mice had higher levels of both ACC1 and FASN (Fig. 6) than did w.t. mice, indicating a potentially higher capacity to synthesize C16:0 by Acsbg1^-/-^ mice. Expression of these enzymes dropped with increasing age in both w.t. and Acsbg1^-/-^ mice. This was particularly evident when the ratio of Acsbg1^-/-^ to w.t. ACC1 was plotted (Fig. 6C), and possibly contributes to the lower VLCFA levels measured in knockout mouse cerebellum. In contrast to ACC1, the malonyl-CoA product of ACC2 is used primarily to regulate FA degradation by the mitochondrial β-oxidation pathway. ACC2 expression levels shown in Fig. 7A did not suggest a significant effect of ACSBG1 deficiency on FA oxidation.

Unlike saturated VLCFA, which can be produced by *de novo* synthesis and subsequent elongation, ω3 and ω6 FA are “essential”, meaning that at least some of these must originate from the diet. However, once ingested, most essential FA can be interconverted [25,35,38]. Interconversion enzymes include desaturases and elongases. Other than in the first few days of life, expression levels of desaturases FADS1 and FADS2 were essentially the same in both w.t. and Acsbg1^-/-^ mice. Expression of ELOVL2 was generally not higher in Acsbg1^-/-^ mice than in w.t. mice, and therefore cannot easily explain the higher level of C20:5ω3 and C22:5ω3 in Acsbg1^-/-^ mice. During the first week of life, ELOVL5 was higher in Acsbg1^-/-^ mice than in w.t., but thereafter was the same or lower, again in contrast to the elevated C20:5ω3 and C22:5ω3 levels. ELOVL1 must be evaluated in future studies, as this enzyme is also required for some interconversion steps in ω3 FA synthesis.

The results of these studies confirm that, despite appealing circumstantial evidence, ACSBG1 does not play a central role in XALD pathophysiology. Findings published by Sheng et al. [31] indicate that absence of ACSBG1 affects testicular Leydig cell function, but any effect(s) on brain, adrenal gland, and ovary remain elusive. Several parameters of cerebellar lipid metabolism are clearly affected when ACSBG1 is defective. Further investigation is needed to clarify these and other roles of ACSBG1 in metabolism.

Metabolic function becomes specialized as development progresses, adapting to tissue-specific needs. Despite its potential role in XALD, the exact function of ACSBG1 remains unclear. Our studies using Acsbg1-/- mice have revealed unexpected findings, particularly regarding the levels of saturated VLCFA in the cerebellum. Contrary to expectations, VLCFA levels were consistently lower in Acsbg1-/- mice, challenging the direct involvement of ACSBG1 in XALD pathology. However, intriguing patterns in monounsaturated and polyunsaturated fatty acids suggest a complex role for ACSBG1 in lipid metabolism and associated disorders. Further research is needed to fully elucidate the function of ACSBG1 and its implications in metabolic diseases.

## ACKNOWLEDGEMENTS

This work was supported by NIH grants NS37355, HD10981, and HD24061

