## Supplementary material for "Role of ACSBG1 in brain lipid metabolism and X-linked adrenoleukodystrophy pathogenesis: Insights from a knockout mouse model": Suppl Fig 1

### Slide 1
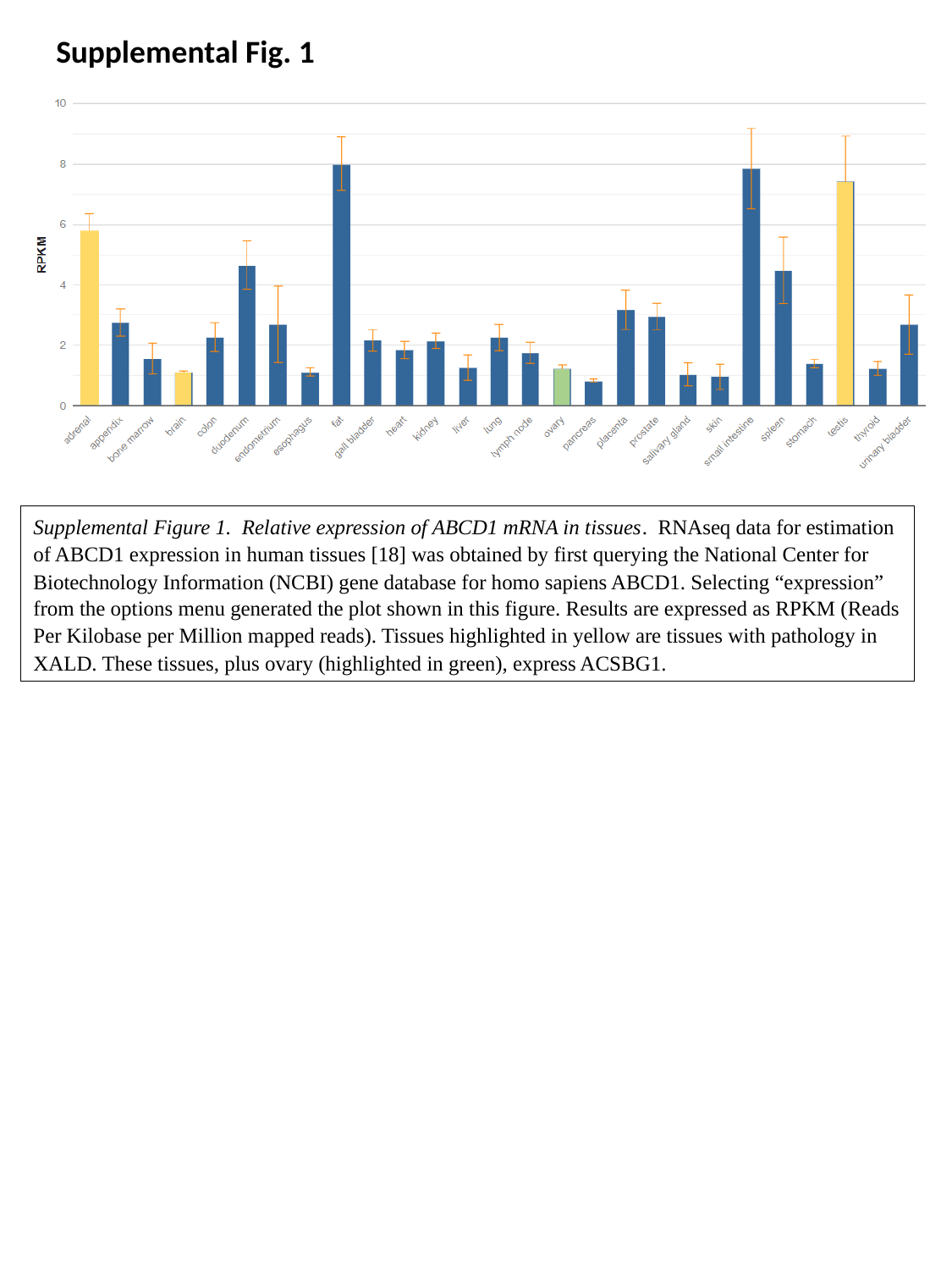

Supplemental Fig. 1
Supplemental Figure 1. Relative expression of ABCD1 mRNA in tissues. RNAseq data for estimation of ABCD1 expression in human tissues [18] was obtained by first querying the National Center for Biotechnology Information (NCBI) gene database for homo sapiens ABCD1. Selecting “expression” from the options menu generated the plot shown in this figure. Results are expressed as RPKM (Reads Per Kilobase per Million mapped reads). Tissues highlighted in yellow are tissues with pathology in XALD. These tissues, plus ovary (highlighted in green), express ACSBG1.
