## Supplementary material for "Role of ACSBG1 in brain lipid metabolism and X-linked adrenoleukodystrophy pathogenesis: Insights from a knockout mouse model": Suppl Fig 2

### Slide 1
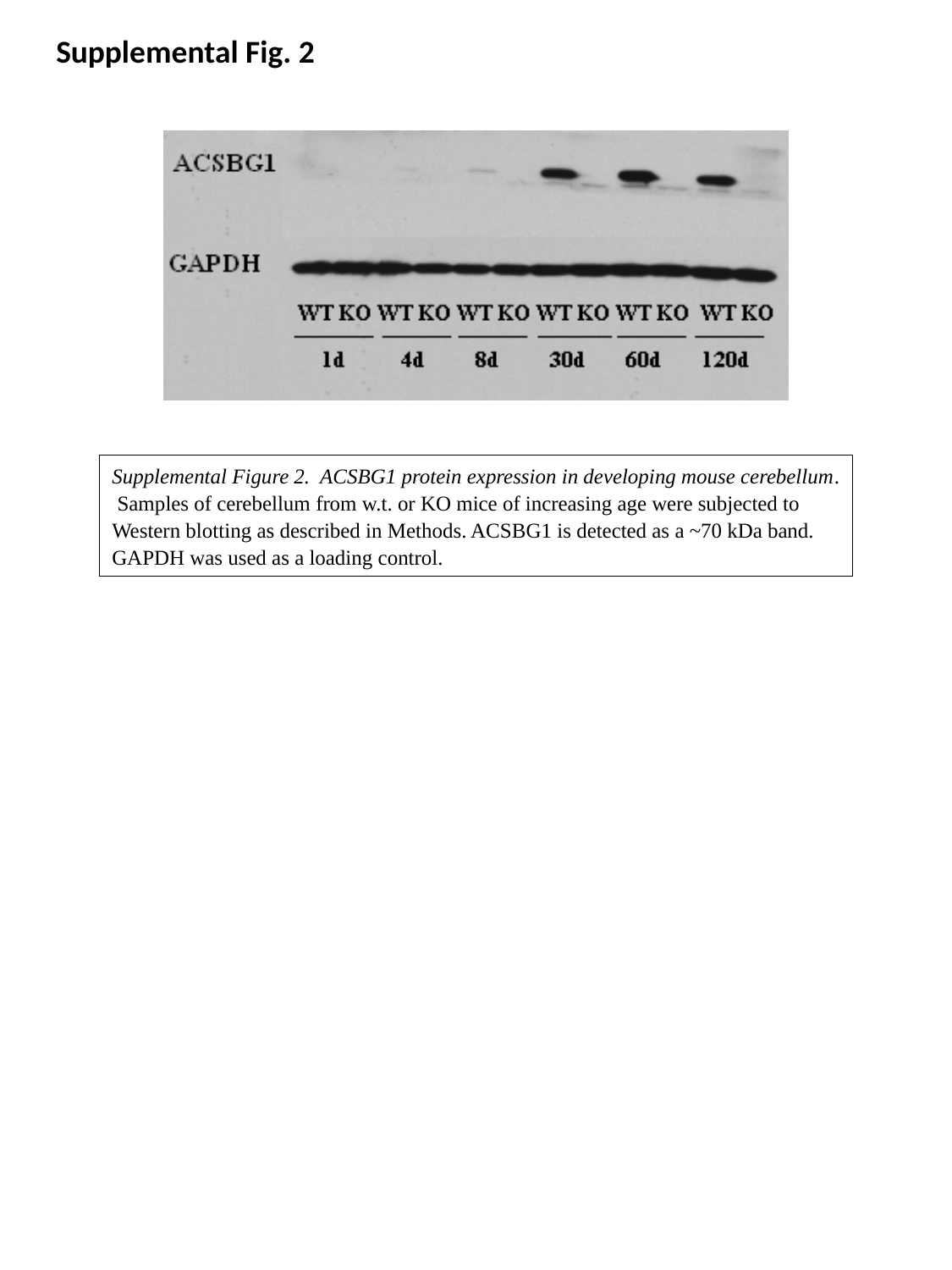

Supplemental Fig. 2
Supplemental Figure 2. ACSBG1 protein expression in developing mouse cerebellum. Samples of cerebellum from w.t. or KO mice of increasing age were subjected to Western blotting as described in Methods. ACSBG1 is detected as a ~70 kDa band. GAPDH was used as a loading control.
