## Supplementary material for "Role of ACSBG1 in brain lipid metabolism and X-linked adrenoleukodystrophy pathogenesis: Insights from a knockout mouse model": Suppl Fig 3

### Slide 1
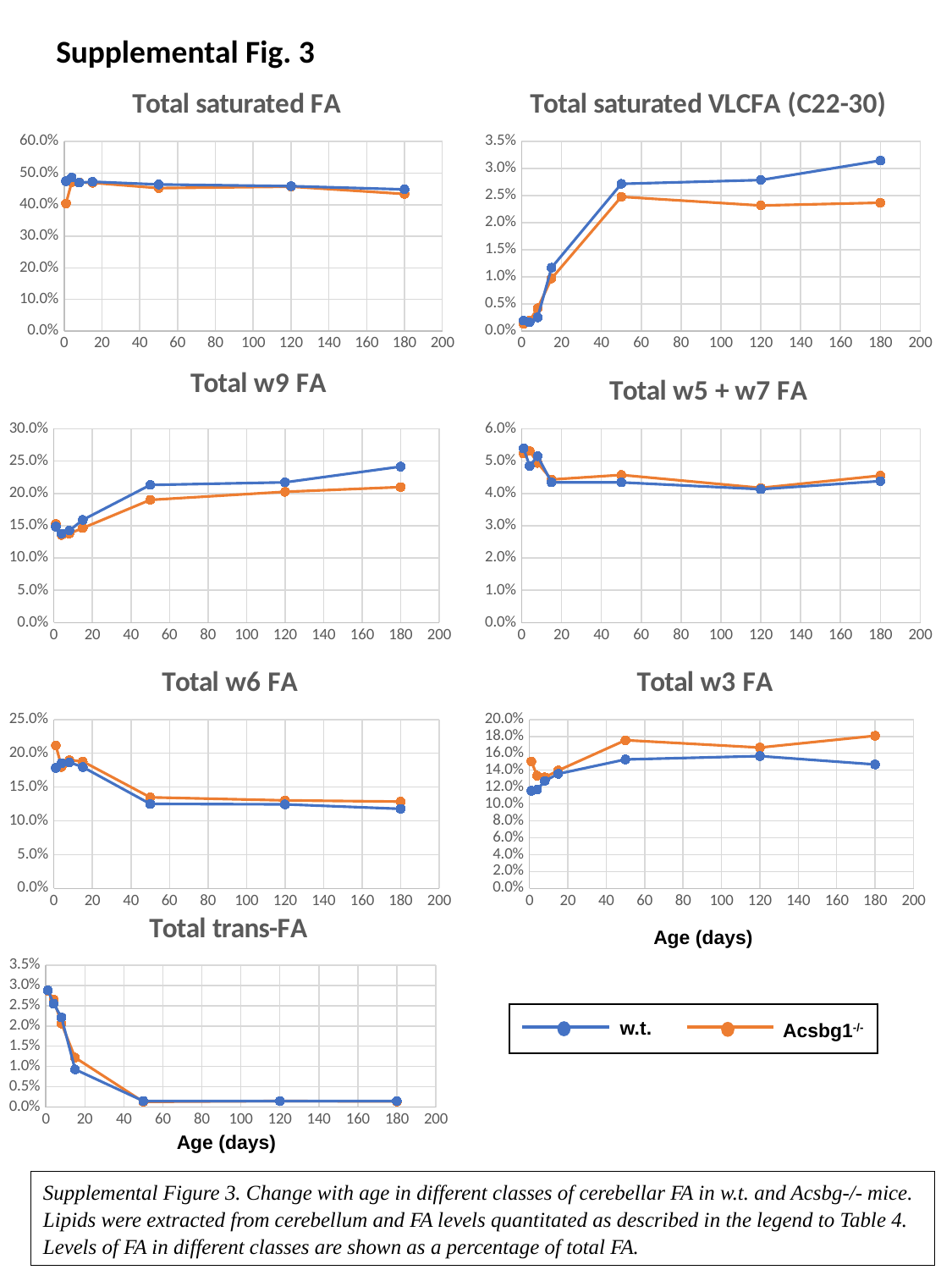

Supplemental Fig. 3
#### Chart: Total saturated FA
| Category | w.t. | Acsbg1-/- |
|---|---|---|
#### Chart: Total saturated VLCFA (C22-30)
| Category | w.t. | Acsbg1-/- |
|---|---|---|
#### Chart: Total w9 FA
| Category | w.t. | Acsbg1-/- |
|---|---|---|
#### Chart: Total w5 + w7 FA
| Category | w.t. | Acsbg1-/- |
|---|---|---|
#### Chart: Total w3 FA
| Category | w.t. | Acsbg1-/- |
|---|---|---|
#### Chart: Total w6 FA
| Category | w.t. | Acsbg1-/- |
|---|---|---|
#### Chart: Total trans-FA
| Category | w.t. | Acsbg1-/- |
|---|---|---|Age (days)
w.t.
Acsbg1^-/-^
Age (days)
Supplemental Figure 3. Change with age in different classes of cerebellar FA in w.t. and Acsbg-/- mice. Lipids were extracted from cerebellum and FA levels quantitated as described in the legend to Table 4. Levels of FA in different classes are shown as a percentage of total FA.
